# Laminin-binding Integrins Regulate Angiogenesis by Distinct and Overlapping Mechanisms in Organotypic Cell Culture Models

**DOI:** 10.1101/2021.06.28.450188

**Authors:** Hao Xu, Susan E. LaFlamme

## Abstract

Endothelial cells engage extracellular matrix and basement membrane components through integrin-mediated adhesion to promote angiogenesis. Our previous studies demonstrated that endothelial expression of laminin-411 and laminin-511 as well as α6 integrins is required for endothelial sprouting and tube formation in organotypic angiogenesis assays. These studies demonstrated that α6 integrins promote migration and regulate the expression of ANGPT2 and CXCR4 and that α6-dependent regulation of CXCR4 contributes to endothelial morphogenesis in our assays. However, these studies did not identify specific roles for the α6β1, α6β4, or α3β1 laminin-binding integrins. Here, we employ RNAi technology to parse the contributions of these integrins. We demonstrate that α6β4 promotes migration, sprouting, and tube formation, and also positively regulates the expression of ANGPT2, but does not promote CXCR4 expression, suggesting that α6β1 functions in this regulation. Additionally, we show that α3β1 regulates endothelial sprouting and tube formation, but is not required for migration in our assays or for the expression of ANGPT2 or CXCR4. Integrin α3β1 promotes the expression of NRP1 and ID1 RNAs, both of which are known to promote angiogenesis. Taken together, our results indicate that laminin-binding integrins play distinct roles during endothelial morphogenesis and do not compensate for one another in organotypic culture.

**Summary Statement:** The laminin-binding integrins α3β1, α6β1, and α6β4 contribute to endothelial sprouting and tube formation in organotypic angiogenesis assays.

## Introduction

Angiogenesis contributes to both normal and pathological processes, including tissue repair, cancer progression, and inflammation (Carmeliet, 2003; Carmeliet, 2005). Angiogenesis is a multistep process that involves the sprouting of new vessels from the preexisting vasculature, which then anastomose to form new vascular networks. Although many mechanisms and regulatory pathways have been identified, a further understanding of the underlying mechanisms that regulate specific aspects of new vessel formation remains an important objective.

At the onset of angiogenesis, endothelial cells interact with proteins present in the extracellular matrix some of which are provided by other cell types, such as those present in the provisional matrix during tissue repair (Eming et al., 2007; Senger and Davis, 2011). Endothelial cells themselves also secrete matrix proteins including fibronectin and the basement membrane components, laminin-411 and laminin-511. The interaction of endothelial cells with these adhesion proteins contributes to the formation and stabilization of endothelial tubes (Avraamides et al., 2008; Hallmann et al., 2020; Hallmann et al., 2005; Senger and Davis, 2011; Song et al., 2017; Turner et al., 2017; Xu et al., 2020).

Endothelial cells express several integrin heterodimers, including the three laminin-binding receptors α3β1, α6β1, and α6β4 (Avraamides et al., 2008; Nishiuchi et al., 2006; Senger and Davis, 2011). Previous studies examined the roles of endothelial laminin-binding integrins during *in vivo* angiogenesis using genetic models demonstrated that these integrins are not required for developmental angiogenesis (Bouvard et al., 2012; da Silva et al., 2010; Dowling et al., 1996; Georges-Labouesse et al., 1996; Germain et al., 2010; Kreidberg et al., 1996; Welser-Alves et al., 2013). More recent studies employed models for the conditional endothelial deletion of either the integrin α3 subunit (Itga3), the α6 subunit (Itga6) or the β4 integrin subunit gene (Itgb4). Integrin α3β1 was found to have an inhibitory effect on pathological angiogenesis (da Silva et al., 2010), whereas α6 integrins were reported to either promote or inhibit angiogenesis depending upon whether α6 alleles were targeted by the expression of either a Tie1-or Tie2-driven Cre recombinase (Bouvard et al., 2012; Bouvard et al., 2014; Germain et al., 2010; Seano et al., 2014). The mechanisms responsible for these disparate phenotypes are not fully understood. However, the phenotype of mice expressing theTie2-driven Cre recombinase exhibited defects due to the loss of the expression of α6 integrins not only in endothelial cells, but also in macrophage and endothelial progenitors (Bouvard et al., 2012; Bouvard et al., 2014; Germain et al., 2010; Seano et al., 2014). It has been difficult to distinguish contributions from α6β1 and α6β4 during angiogenesis, as the integrin β1 subunit dimerizes with multiple α subunits in endothelial cells making it difficult to discern functions specific to α6β1. Mouse genetic studies examining the effect of Tie2-dependent deletion of the β4 subunit identified roles for α6β4 in vessel remodeling and endothelial barrier function in the brain vasculature (Welser et al., 2017; Welser-Alves et al., 2013). However, the contribution of endothelial specific expression of α6β4 to early aspects of angiogenesis has not been explored.

To dissect the roles of individual laminin-binding integrins in early stages of angiogenesis, we employed RNAi technology together with two organotypic assays, which model angiogenesis in an ECM environment similar to that present during tissue repair (Bajaj et al., 2012; Bishop et al., 1999; Nakatsu and Hughes, 2008). Using these assays, we previously demonstrated that depleting endothelial cells of α6 integrins inhibited endothelial sprouting and tube formation (Xu et al., 2020). We showed that α6 integrins promoted migration, the expression of their substrate, laminin-511, as well as the angiogenesis-associated genes angiopoitin-2 (ANGPT2) and the chemokine receptor, CXCR4. We further demonstrated that recombinant CXCR4 partially rescued endothelial morphogenesis when the expression of α6 integrins was inhibited (Xu et al., 2020). However, our previous studies did not distinguish contributions from α6β1 or α6β4 or whether the α3β1 laminin-binding integrin also contributes endothelial morphogenesis in these models. Our new results indicate that the depletion of α6β4 with three distinct siRNAs targeting the β4 subunit resulted in the inhibition of migration, as well as endothelial sprouting and tube formation. Depletion of α6β4 also reduced the expression of the angiogenesis-associated genes, ANGPT2 and neuropilin-1 (NRP1), but did not inhibit the expression of CXCR4, suggesting that CXCR4 expression may be α6β1-dependent. RNAi-mediated depletion of α3β1 by targeting the integrin α3 subunit also inhibited endothelial morphogenesis. However, the depletion of α3β1 did not affect migration or the expression of ANGPT2, CXCR4 or laminin-511. Interestingly, the expression of the angiogenesis-associated genes, neurophilin-1 (NRP1) and the inhibitor of DNA binding -1 (ID1) was significantly decreased in α3-depleted endothelial cells, suggesting a potential mechanism for the contribution of α3β1 in our assays. Taken together, these results suggest that α6β1, α6β4 and α3β1 can impact endothelial morphogenesis by distinct and non-redundant mechanisms.

## Results

### Integrin α6β4 is expressed on veins and small vessels of the dermis

Previous studies showed the co-expression of α6β4 and its ligand laminin-511 in tumor-associated angiogenic vessels (Nikolopoulos et al., 2004). Others have reported that α6β4 expression is restricted to arterioles (Welser-Alves et al., 2013). Interestingly, single-cell RNA sequencing data indicated restricted expression for β4 mRNA in the brain vasculature; however, the same study found that in the lung vasculature that β4 mRNA was widely expressed in endothelial cells, including veins and venules (He et al., 2018). Since the β4 subunit only dimerizes with the α6 subunit (Sonnenberg et al., 1990), these results suggest that the endothelial expression of α6β4may be tissue specific and its restriction to arterioles may be specific for the brain vasculature. To confirm this at the protein level, we examined α6β4 expression by immunofluorescence microscopy. In the adult murine retinal vasculature, α6β4 expression colocalized with the strong expression of α-smooth muscle actin (α-SMA), consistent with its expression being restricted to arteries/arterioles in the brain (Fig. S1). However, in the dermal vasculature, α6β4 expression is not restricted by α-SMA-positive arterioles (Fig. 1) and α6β4 is not expressed by lymphatic vessels (Fig. S2), indicating α6β4 is expressed in veins and small vessels including venules. This expression pattern in the dermal vasculature suggests that α6β4 can contribute to the regulation of angiogenesis.

**Figure 1.**
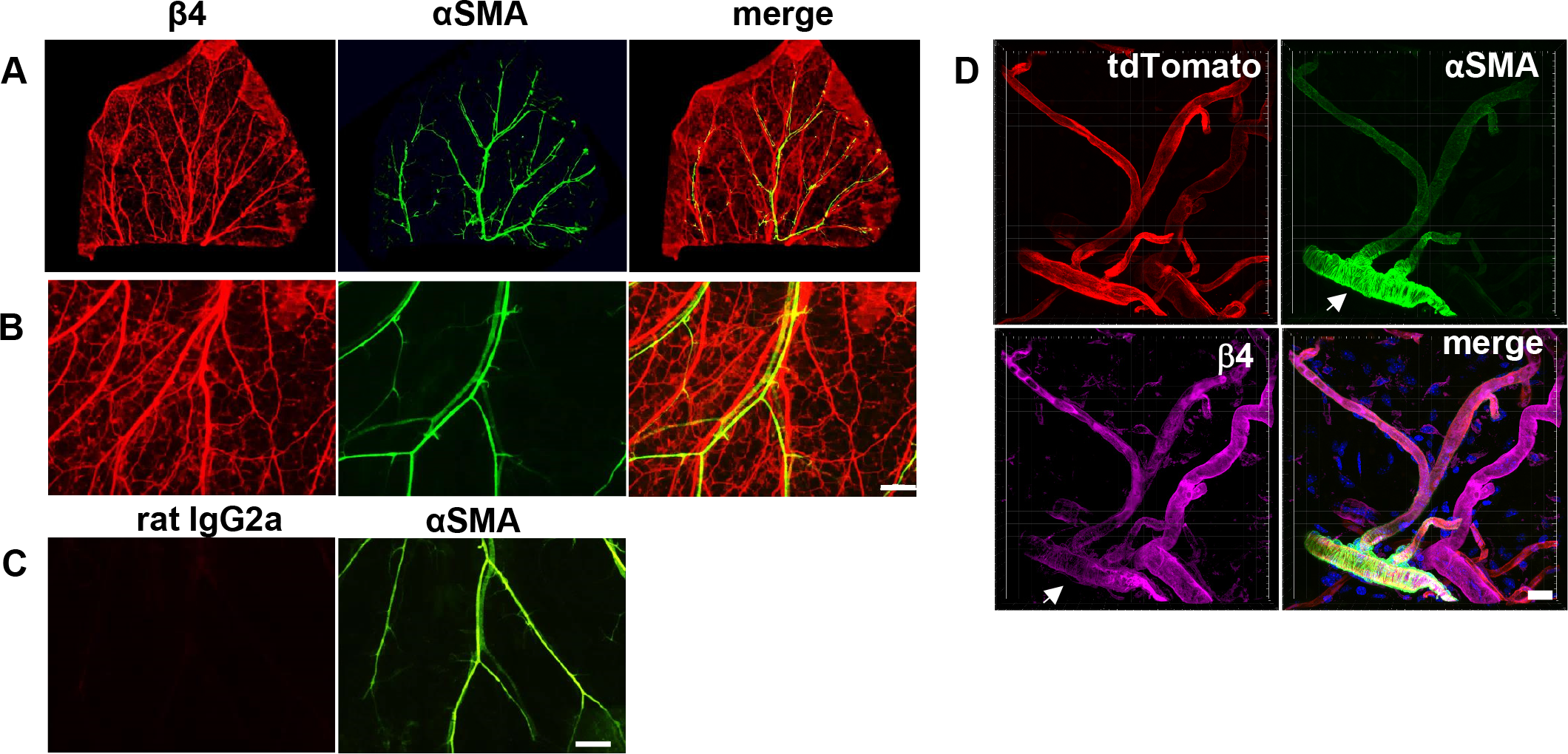
The α6β4 integrin is expressed in veins and venules of the dermal vasculature. **(A)** Immunofluorescence staining of whole mount mouse ear for α-smooth muscle actin (α-SMA) and β4-integrin subunit. **(B)** Image of β4 and α-SMA expression acquired at a higher magnification. Scale = 500 μm. (C) Rat IgG2a control for staining with the antibody to the β4 subunit. **(D)** Confocal immunofluorescence staining of mouse skin for α-smooth muscle actin and β4-integrin subunit. Vasculature is identified by tdTomato (red) expression. Image is a maximum projection of acquired z-stack. Arrowhead indicates the location of an arteriole. Scale = 20 μm.

### Integrin α6β4 promotes endothelial migration

Using HUVECs, we previously demonstrated that α6 integrins promote endothelial morphogenesis, in part by promoting cell migration, as well as the expression of CXCR4, ANGPT2, and the α5 chain (LAMA5) of laminin-511 (Xu et al., 2020). However, we did not determine the specific contributions of α6β1 and α6β4. Since HUVECs express α6β4 on their cell surface (Fig, S3), we used the same organotypic co-culture approaches to determine the contribution of α6β4. To inhibit the expression of α6β4, we were able to use an RNAi approach to target the integrin β4 subunit, because the β4 subunit only dimerizes with the α6 subunit (Sonnenberg et al., 1990). We depleted the β4 subunit using three distinct siRNA targeting sequences in three independent experiments and measured the affect in Transwell migration assays. The efficiency of knockdown was determined by qPCR and western blot (Fig. 2A-B). Since cell motility is an important aspect on angiogenesis, we examined the role of α6β4 in endothelial cell migration. The depletion of β4 resulted in a significant reduction of cell migration across gelatin-coated filters (Fig. 2C). Importantly, this phenotype was consistent across all three siRNA targeting sequences and consistent with previous studies that suggested a role for α6β4 in the regulation of endothelial cell migration (Nikolopoulos et al., 2004).

**Figure 2.**
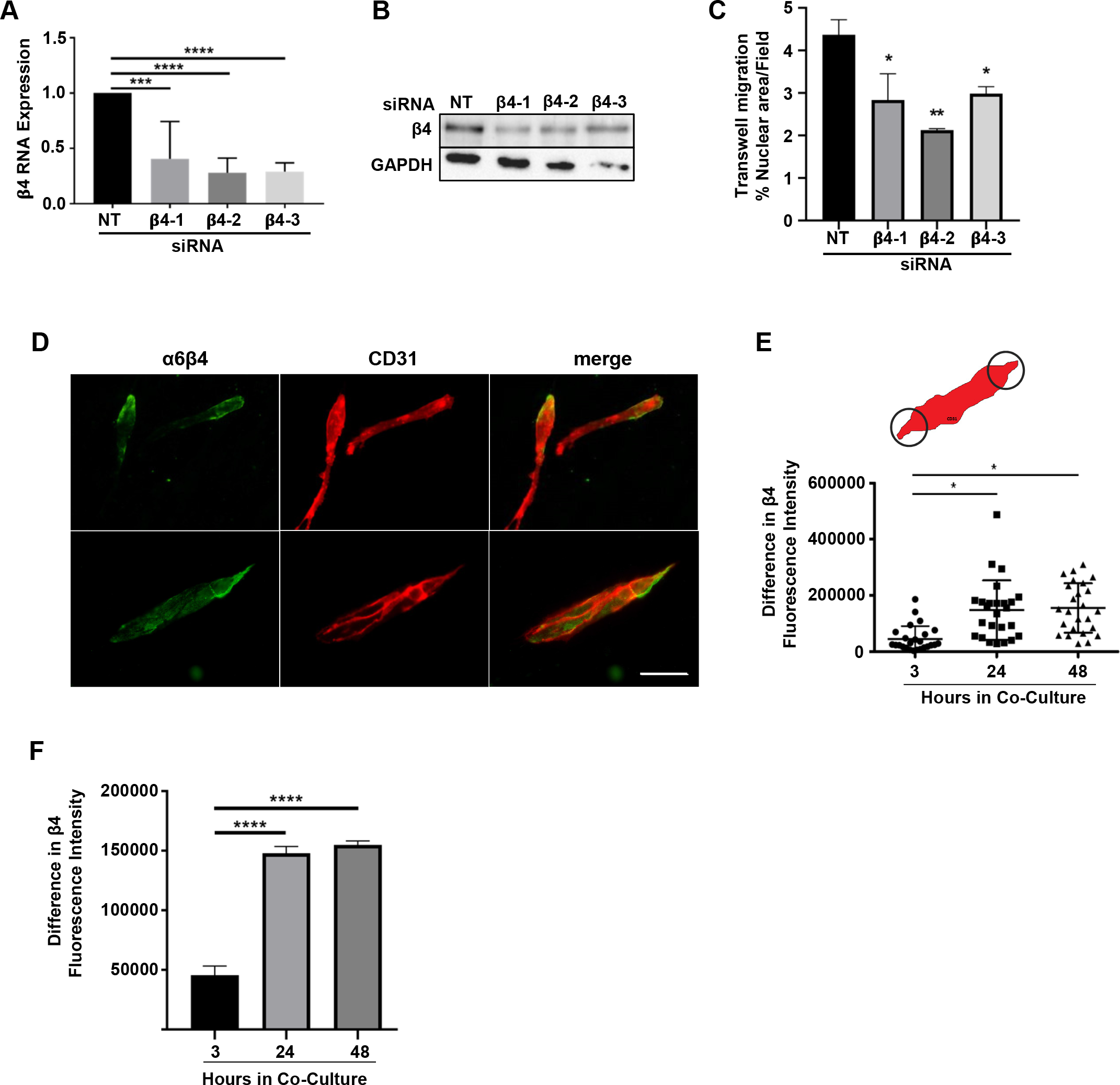
The expression of α6β4 promotes cell migration and becomes polarized on forming endothelial cords. **(A)** The efficiency of β4 depletion using 3 siRNA targeting sequences was determined by qPCR. Data were normalized to β-actin and then to non-targeting (NT) control. Plotted is the mean RNA expression ± s.d. n = 5 independent experiments. **(B)** Representative western blot showing the efficiency of β4 depletion at the protein level. **(C)** Transwell migration assays with non-targeting (NT) or β4-targeting siRNA treated cells. Fifteen fields were analyzed from each of two independent experiments. Plotted is the mean percentage of area per field stained for nuclei ± s.e.m. n = 2 independent experiments. Data were analyzed by one-way ANOVA and Dunnett’s multiple comparisons test. **(D-F)** The polarized expression of β4 in endothelial cells was analyzed at 3h, 24h, and 48 h. **(D)** Representative images of endothelial cells at the 24 h immunostained for the β4 integrin subunit (green) and CD31 (red). Scale = 100 µm. **(E)** Plotted is the mean difference in β4/CD31 fluorescence intensity (at 3, 24, and 48 h)) between polar ends of endothelial cells or cell cords using a constant ROI (See top panel in E for diagram) from three independent experiments ± s.d. n = 25 cells. **(F)** The same data shown in Panel E plotted as the mean difference in β4/CD31 fluorescence intensity between polar ends of endothelial cells or cell cords using a constant ROI ± s.e.m. n = 3 independent experiments. Data were analyzed by one-way ANOVA and the Tukey’s multiple comparisons test.*p ≤ 0.05, **p ≤ 0.01, ***p ≤ 0.001, ****p ≤ 0.0001.

### Integrin α6β4 expression is polarized during the formation of endothelial tubes

Previous studies described the localization of α6β4 at the tips of sprouting vessels *in vivo* (Enenstein and Kramer, 1994). Thus, we were interested to examine the localization of α6β4 during the formation of endothelial tubes in organotypic assays. In our planar co-culture assays, endothelial cells attach and spread within 1 h and begin to elongate, migrate and form cords after approximately 24 h (Mavria et al., 2006; Xu et al., 2020). Therefore, we employed immunofluorescence microscopy to examine the localization of endothelial α6β4 distribution at 3 h, 24 h and 48 h by immunofluorescence microscopy. Co-cultures were immunostained for the β4 subunit and the endothelial marker, CD31. This was feasible as fibroblasts that are present in the co-culture do not express either α6β4 or CD31. To measure the polarized distribution of α6β4 in endothelial cells/cords, we used CD31 to establish the endothelial area and determined the difference in β4 fluorescence intensity on either ends of endothelial cells or cords using a constant-sized ROI (Fig. 2D-F). At 3 h of co-culture, individual endothelial cells are easily distinguishable. Little polarized localization was observed at this time (Fig. 2E-F). At 24 and 48 h when endothelial cords had begun to form cords (Mavria et al., 2006; Xu et al., 2020), the localization of α6β4 was predominately stronger at one end of the endothelial structure (Fig. 2E-F). Although we cannot discern whether the concentrated localization occurs at the migrating front, the polarized distribution of α6β4 is consistent with its role in endothelial migration and the localization of α6β4 in sprouting vessels *in vivo*.

### Integrin α6β4 promotes endothelial morphogenesis

The contribution that endothelial integrin α6β4 makes to early stages of angiogenesis in the adult has not yet been examined. To determine whether endothelial α6β4 contributed to the defective sprouting phenotype that we previously observed for α6-depleted endothelial cells (Xu et al., 2020), we analyzed 6-day sprouts of endothelial cells depleted of β4 compared to control. The efficiency of β4 depletion with three targeting siRNAs is shown in Fig. 1A, as the same batch of control and β4-depleted endothelial cells were used in studies to examine morphogenesis. Shown are representative images of bead-sprout assays in which endothelial cells were depleted of β4 with each of the three β4 targeting siRNAs compared to control (Fig. 3A). Knockdown with each of these three sequences resulted in a decrease in both sprout length (Fig. 3B and D), as well as the number of sprouts per bead (Fig. 3C and E).

**Figure 3.**
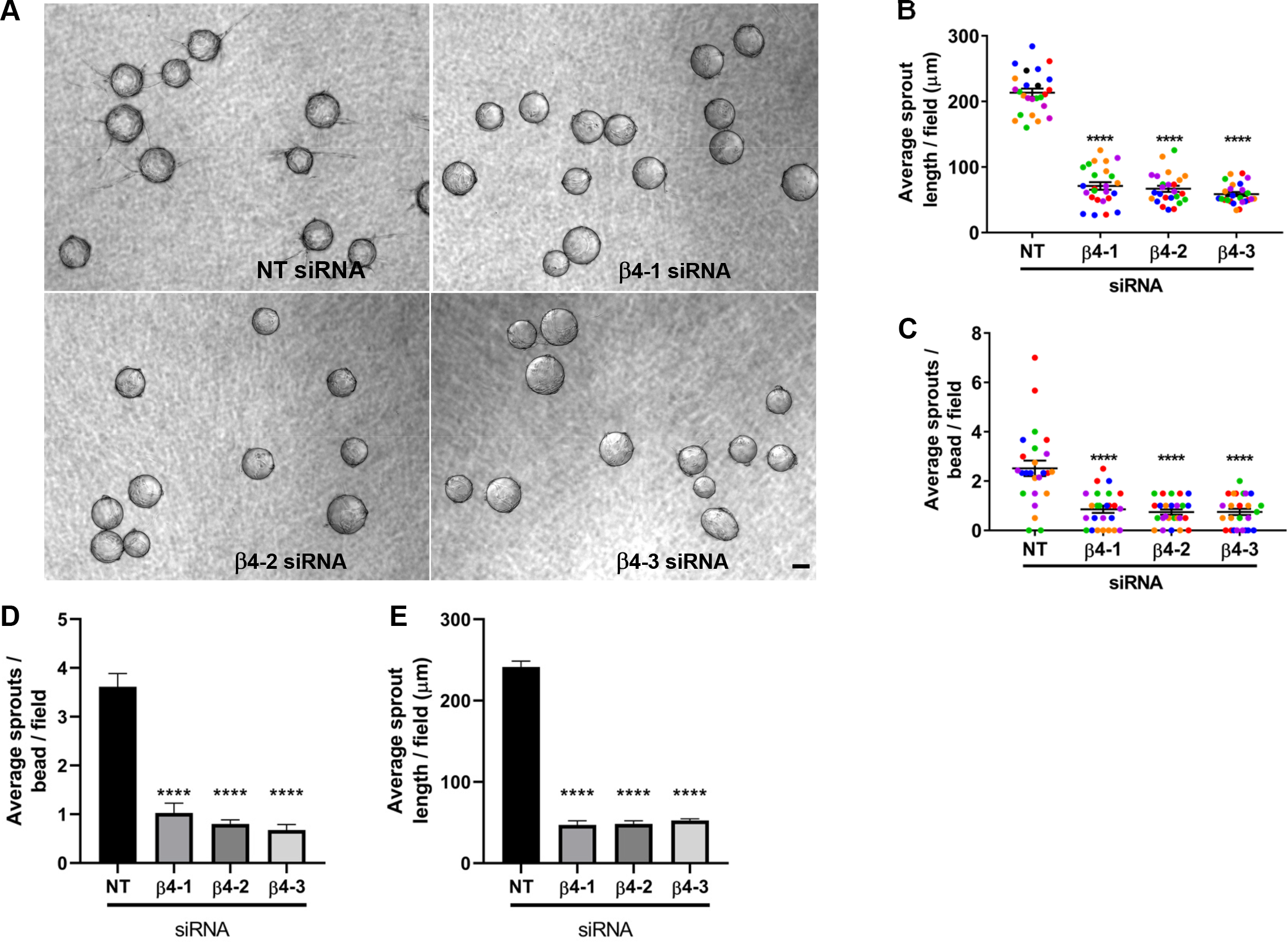
Integrin α6β4 regulates endothelial morphogenesis. Endothelial cells from the same batches of endothelial cells depleted of the β4 subunit used in experiments presented in Fig. 2 were employed in bead sprout assays. **(A)** Shown are representative images of 6-day bead sprouts transfected with non-targeting and β4-targeting siRNA sequences 1-3. Scale = 100 μm. **(B-C)** Depletion of α6β4 expression inhibits sprouting. Quantitation of sprout length (top) and number of sprouts (bottom) from 6-8 beads in each of 5 randomly selected fields in 5 independent experiments. **(B)** Plotted is the average sprout length per field ± s.e.m n = 25 fields. **(C)** Quantitation of the average sprouts per bead per field ± s.e.m. Each independent experiment is represented by a distinct color. **(D-E)** The average sprout length **(D)** and average sprouts per bead **(E)** was calculated for each independent experiment and the data plotted as the mean ± s.e.m. n= 5 independent experiments. Data were analyzed using two-tailed Student’s t-test. Each independent experiment is represented by a distinct color. ***p ≤ 0.001, ****p ≤ 0.0001.

### Integrin a6β4 promotes the expression of ANGPT2 and other angiogenesis associated genes

Since we determined that α6β4 contributed to both cell migration and endothelial morphogenesis, we next sought to determine whether it also contributed to the regulation of the angiogenesis-associated genes that we previously identified to be inhibited in α6-depleted cells. Expression of LAMA5, CXCR4, and ANGPT2 RNA transcripts were previously determined to be positively associated with the expression of endothelial α6 integrins (Xu et al., 2020). To evaluate the contribution of α6β4 to the expression of the genes, we isolated RNA from endothelial cells treated with non-targeting and β4-targeting siRNA from 6-day bead sprout assays shown in Figure 3 and analyzed gene expression by qPCR (Fig. 4). Interestingly, a significant decrease in ANGPT2 expression was observed with all three siRNA targeting sequences (Fig. 4B), indicating that α6β4 contributes to the regulation of ANGPT2 RNA expression. A significant decrease in the expression of the α5 chain (LAMA5) of laminin-511 was apparent with two siRNA targeting sequences (Fig. 4C) with the of expression of LAMA5 in cells targeted with the third siRNA trending downward. These data suggest that α6β4 is likely to also contributes to the expression of LAMA5 RNA. Notably, the expression of CXCR4 was not inhibited in β4-depleted endothelial cells (Fig. 4D), suggesting that α6β1 is likely to be responsible for the regulation of CXCR4 by α6 integrins. Similar to our published results for α6-depleted endothelial cells, the expression of LAMA4 was not affected in β4-depleted cells (Fig. 4E).

**Figure 4.**
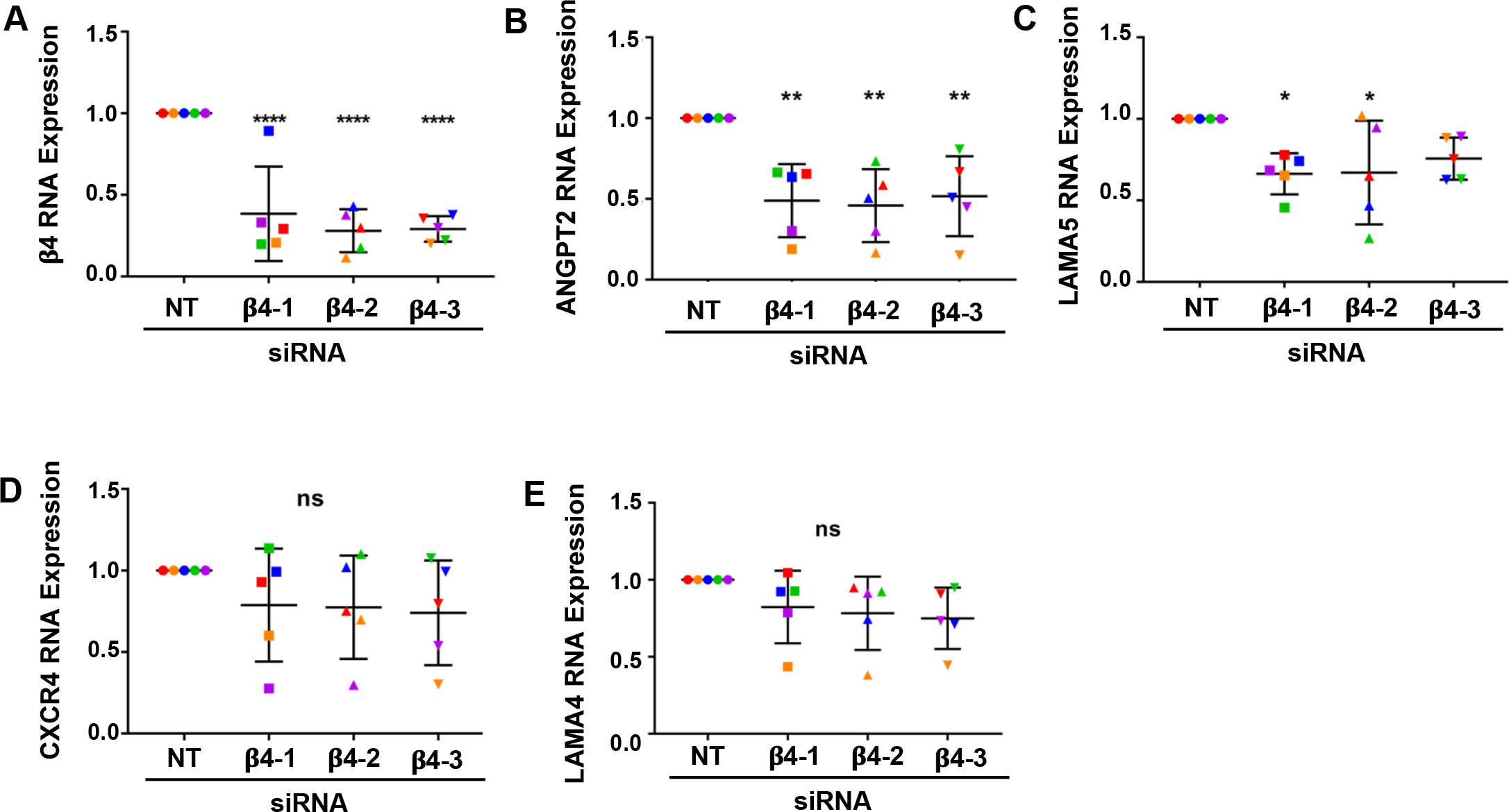
The expression ANGPT2 is positively regulated by integrin α6β4. RNA was isolated from 6-day bead sprouts and analyzed by qPCR. **(A)** Shown is the efficiency of β4 depletion using 3 siRNA targeting sequences (also shown in Fig. 1A) and the effects of β4 depletion on the expression of the **(B)** ANGPT2, **(C)** LAMA5, **(D)** CXCR4, and **(E)** LAMA4. Data were normalized to β-actin and then to the non-targeting (NT) control. Plotted is the mean RNA expression ± s.d. n = 5 independent experiments. Data were analyzed by one-way ANOVA and Dunnett’s multiple comparisons test. Each independent experiment is represented by a different color. ns = not significant, *p ≤ 0.05, **p ≤ 0.01, ****p ≤ 0.0001.

To determine whether the depletion of α6β4 integrin affected the expression of other angiogenesis-associated genes (De Smet et al., 2009; del Toro et al., 2010; Strasser et al., 2010), we analyzed the expression of DLL4, JAG1, JAG2, KDR, NRP1 ID1, ID2, and PDGFB genes by qPCR (Fig. 5A-C). The results indicate that the expression of NRP1 RNA was significantly inhibited by all three siRNAs. Additionally, a significant downregulation of PDGFB RNA was observed with two of the three β4-targeting siRNAs with the third siRNA exhibiting a downward trend (Fig. 5A-B). Taken together, these results suggest that α6β4 contributes to positive regulation of the expression of ANGPT2 RNA and NRP1 RNAs and likely to the expression of PDGFB and LAMA5 RNAs as well.

**Figure 5.**
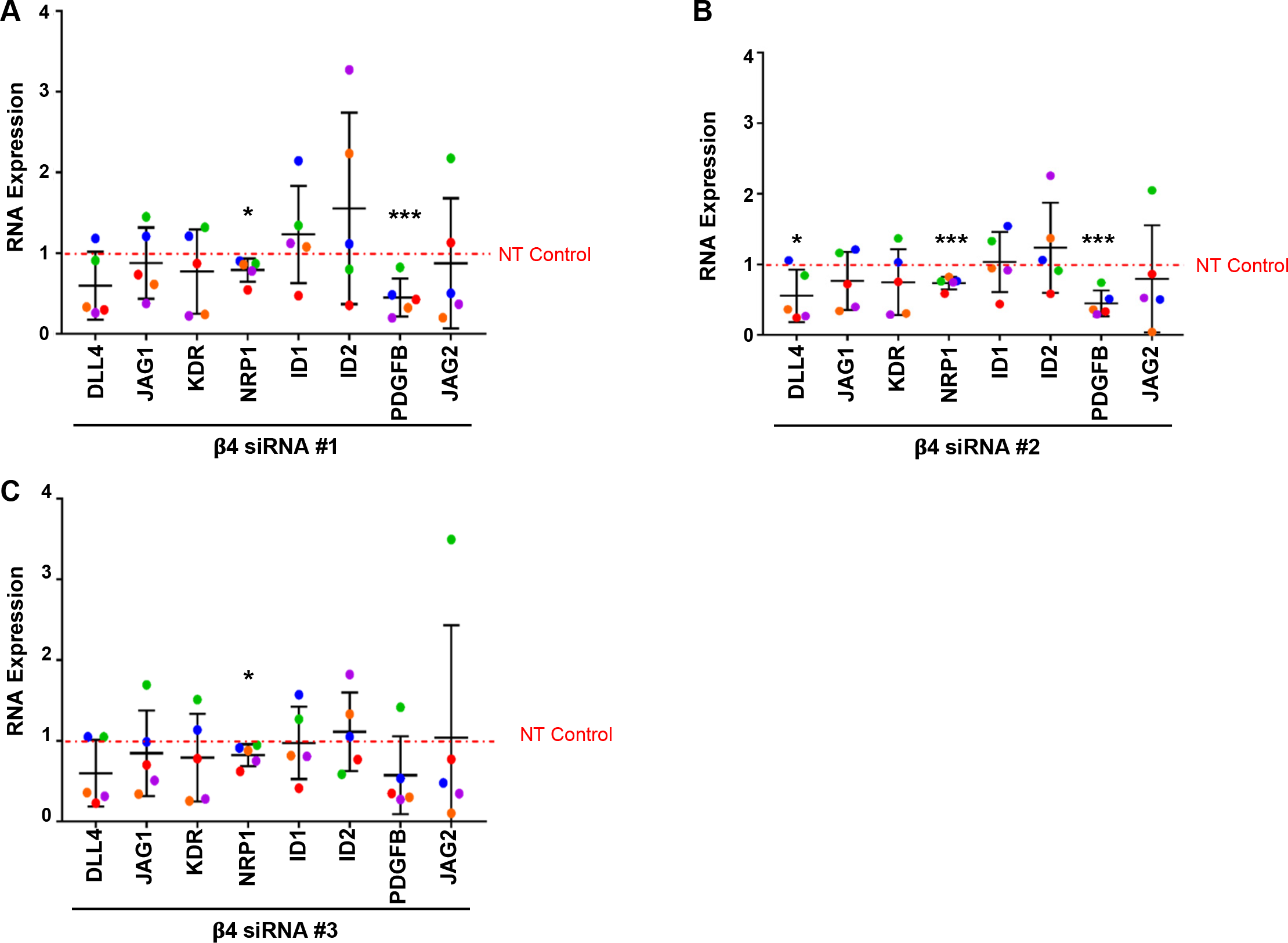
The expression of NRP1 and PDGFB are positively associated with integrin α6β4 expression. Changes in RNA expression in 5-day bead sprout assays of β4-depleted endothelial cells and control cells were measured by qPCR. The efficiency of β4 depletion using 3 siRNA targeting sequences was previously shown in Fig. 4. Data were normalized to β-actin and then to non-targeting (NT) control. Plotted is the mean RNA expression ± s.d. n = 5 independent experiments. Mean expression of each gene was compared to non-targeting (NT) control and analyzed by two-tailed Student’s t-test. *p ≤ 0.05, ***p ≤ 0.001.

### Integrin α3β1 plays overlapping and distinct roles during endothelial morphogenesis

In addition to α6β1 and α6β4, endothelial cells also express integrin α3β1 to engage their laminin substrates (Nishiuchi et al., 2006; Stenzel et al., 2011). Since the expression of α3β1 was found to be a negative regulator of angiogenesis *in vivo* (da Silva et al., 2010), we questioned whether it served a similar role in our organotypic cultures. To determine whether depletion of α3β1 enhanced endothelial morphogenesis, we employed lentiviral vectors for the doxycycline inducible expression of either non-targeting (NT) or α3-targeting shRNA that was accompanied by the expression of an RFP reporter. Surprisingly, the efficient depletion of the α3 subunit from endothelial cells inhibited both cord formation in planar co-cultures, as well as sprouting in bead-sprout assays (Fig. 6A-C). Images of 6-day planar co-cultures and bead-sprout assays with endothelial cells expressing either NT or α3-targeting shRNA together with RFP are shown in Fig. 6 E-F. It is noteworthy that sprouting was severely inhibited in α3-depleted cells. However, unlike the depletion of α6-integrins, depletion of α3β1 did not impact cell migration across gelatin-coated Transwells (Fig. 6D).

**Figure 6.**
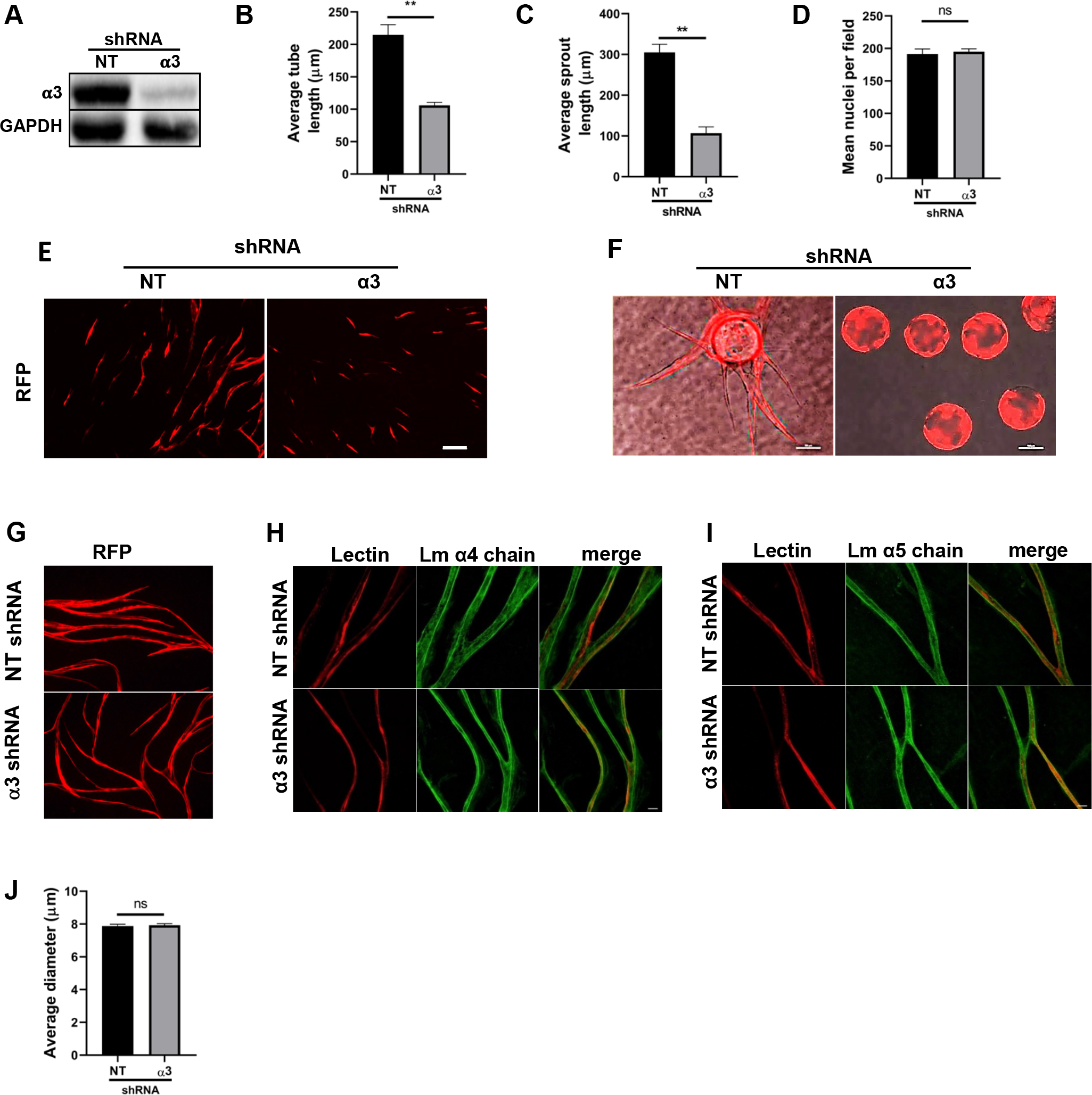
Depletion of endothelial integrin α3β1 inhibits early endothelial tubular morphogenesis but does not disrupt established tube morphology. **(A)** Representative western blot of the expression of the α3 integrin subunit after induction of α3-targeting, or non-targeting (NT) shRNA. **(B)** Average length of tubes/field from each experiment were calculated and the data plotted as the mean ± s.e.m. n= 3 independent experiments **(C)** Plotted is the mean sprout length measured from 3 independent experiments of day 6 bead sprout assays with 10 randomly selected fields each, averaging 6-8 beads per field ± s.e.m n = 3 independent experiments. **(D)** Analysis of nuclear staining in transwell migration assays with non-targeting (NT) or α3-targeting cells. Five fields were analyzed from each of three independent experiments. Plotted is the mean area of nuclei per field, normalized to number of cells seeded ± s.e.m. n= 3 independent experiments. **(E)** Representative images of day 6 planar co-culture. Shown are endothelial cells expressing RFP in conjunction with shRNA under doxycycline regulation. Scale = 250 µm. **(F)** Representative images of day 6 bead sprout assay. Phase/Fluorescent image overlays of endothelial cells expressing RFP in conjunction with shRNA under doxycycline regulation. Scale = 100 µm. Data were analyzed by two-tailed Student’s t-test. **(G - J)** Endothelial tubes were formed for 10 days in the planar co-culture assay and the expression of α3- or non-targeting (NT) shRNA was induced with doxycycline for 12 days. **(G)** Expression of RFP fluorescence **(G)**, laminin-411 **(H)** and laminin-511 **(I)** was analyzed after 12 days of induction of either α3- or non-targeting (NT) shRNA. Scale = 25 µm. **(J)** The average tube width per field in (NT-sh) and (α3-sh) structures measured from ten randomly selected fields in three independent experiments. The mean tube width ± s.e.m. n = 3 independent experiments. Data were analyzed by two-tailed Student’s t-test. ns = not significant, ****p ≤ 0.0001.

Since the depletion of α6-integrins from established endothelial tubes resulted in defective tube morphology and the striking loss of laminin-511 from established tubes (Xu et al., 2020), we asked whether α3β1 also contributed to maintenance of established endothelial tube morphology. Endothelial cells transduced with either α3-targeting or non-targeting (NT) inducible-shRNA were placed in planar-co-cultures for 10 days in the absence of doxycycline to allow for the establishment of endothelial tubes. Expression of shRNAs was then induced with doxycycline for 12 days as indicated by the expression of RFP (Fig. 6G). Co-cultures were then fixed and immunostained for either the laminin-α4 or the laminin-α5 chains along with TRITC-lectin for enhanced endothelial cell visualization. (Fig. 6H-I). Fluorescence imaging revealed that both laminin-411 and laminin-511 are present and are strongly associated with the endothelial tubes, which maintained their morphology after the induction of either NT or α3-targeting shRNA (Fig. 6H-J). Thus, unlike α6 integrins, α3β1 appears to be dispensable for the maintenance of the morphology of established endothelial tubes and their association with laminins at least in the context of our organotypic assays.

### Integrin α3β1 contributes to the expression of a distinct set of angiogenesis-associated genes

Since depletion of α3β1 from endothelial cells shared a subset of phenotypes with α6-depleted cells, we asked whether any of the genes that are regulated by α6 integrins are also regulated by α3β1. Depletion of the α3 integrin subunit was achieved using siRNAs with three distinct targeting sequences and efficiency determined by qPCR (Fig. 7A). Similar to the phenotype observed when α3 was depleted by shRNA, depletion with all three siRNAs resulted in the inhibition of endothelial morphogenesis (Fig. 7B). Interestingly, qPCR analysis of RNA harvested from α3-depleted endothelial cells from 6-day bead-sprout assays did not reveal significant changes in LAMA4, LAMA5, ANGPT2 or CXCR4 expression (Fig. 7C-F). However, expanded qPCR analysis for other angiogenesis-associated genes identified a significant downregulation of NRP1 and ID1 with all three siRNA targeting sequences and PDGFB with two targeting sequences the third showing a downward trend (Fig. 8A-C). These results suggest that α3β1 regulates endothelial morphogenesis by distinct mechanisms and that the loss of its expression cannot be compensated by α6β1 or α6β4.

**Figure 7.**
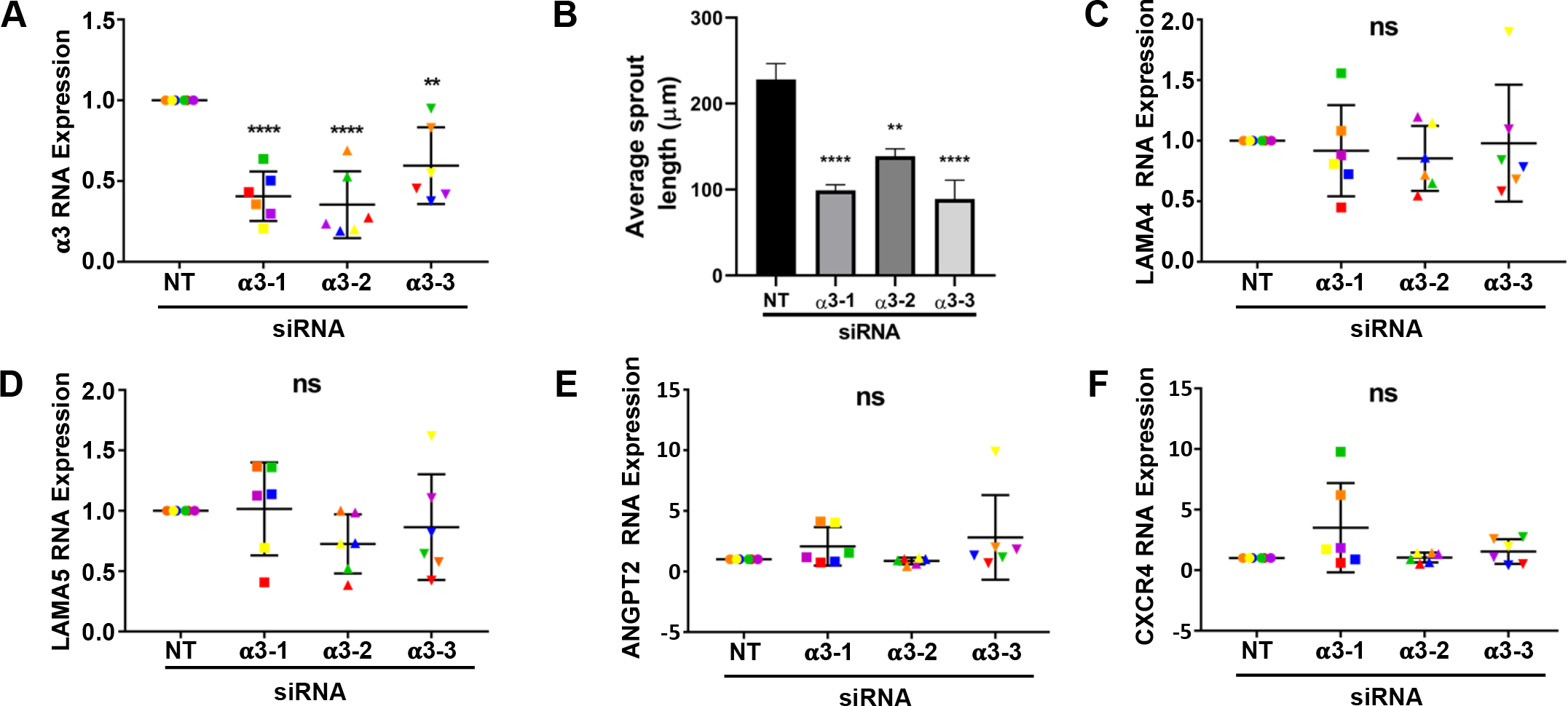
Integrin α3β1 expression is not required for the expression of CXCR4, ANGPT2, LAMA4, and LAMA5. **(A)** The efficiency of α3 depletion using 3 siRNA targeting sequences was determined by qPCR from 6 independent experiments. Data were normalized to β-actin and then to non-targeting (NT) control. Plotted is the mean RNA expression ± s.d. n = 6 independent experiments**. (B)** Quantitation of sprout length from 6-8 beads in each of 5 randomly selected fields in 6 independent experiments. Data plotted as the mean sprout length ± s.e.m. n= 6 independent experiments. **(C-F)** The effects of α3 depletion on the expression of the LAMA5, LAMA4, ANGPT2, and CXCR4. Data were normalized to β-actin and then to non-targeting (NT) control. Plotted is the mean RNA expression ± s.d. n = 6 independent experiments. Data were analyzed by one-way ANOVA with Dunnett’s multiple comparisons test. Each independent experiment is represented by a distinct color. ns = not significant, **p ≤ 0.01, ****p ≤ 0.0001.

**Figure 8.**
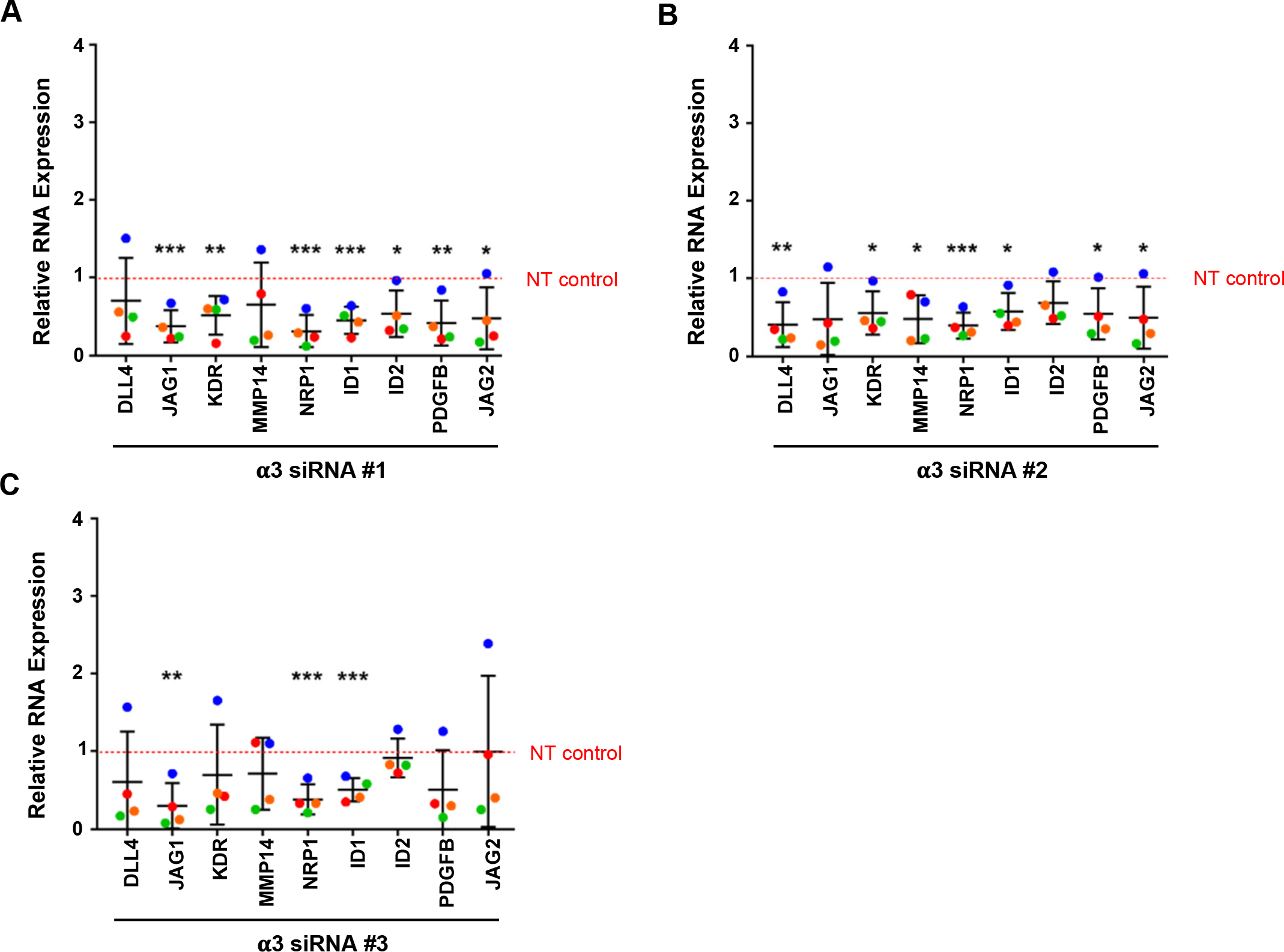
Regulation of angiogenesis-associated genes by integrin α3β1. Changes in RNA expression in 5-day bead sprout assays of α3-depleted endothelial cells and control were measured by qPCR. The efficiency of α3 depletion using 3 siRNA targeting sequences previously shown in Fig. 6. Data were normalized to β-actin and then to non-targeting (NT) control. Plotted is the mean RNA expression ± s.d. n = 4 independent experiments. Mean expression of each gene was compared to non-targeting (NT) control and analyzed by two-tailed Student’s t-test. *p ≤ 0.05, **p ≤ 0.01, ***p ≤ 0.001.

## Discussion

Taken together, our data suggest that the endothelial laminin-binding integrins play overlapping and distinct roles during endothelial morphogenesis (Fig. 9). Our previous studies indicated that expression of integrin α3β1 does not compensate for the depletion of α6 integrins in our assays (Xu et al., 2020). Our current findings demonstrate that inhibiting the expression of α3β1 also inhibits morphogenesis, indicating that α6 integrins do not compensate for the loss of α3β1. Because the integrin β1 subunit dimerizes with multiple α subunits, we were unable to directly identify specific roles for α6β1. However, we did demonstrate that the α6β4 integrin plays an essential role in promoting both endothelial sprouting and tube formation, suggesting that both α6β1 and α6β4 may contribute to these processes. Transcriptome analysis indicates that α3β1, α6β1 and α6β4 regulate distinct sets of angiogenesis-associated genes at the level of RNA expression (see Supplementary Table 4). For example, our previous studies indicated that depletion of the α6 subunit inhibited the expression of both ANGPT2 and CXCR4; however, we did not identify specific contributions of α6β1 or α6β4 (Xu et al., 2020). Here we show that depletion of α6β4 resulted in the inhibition of the expression of ANGPT2 RNA; however, the downregulation of α6β4 did not alter the expression of CXCR4. Taken together these data suggest that α6β1 is the α6 integrin responsible for regulating the expression of CXCR4 and that α6β1 regulates endothelial morphogenesis at least in part without contribution from α6β4, as the expression of recombinant CXCR4 partially rescued endothelial morphogenesis when the α6 subunit was depleted (Xu et al., 2020). Notably, depletion of α3β1 did not alter the expression of ANGPT2 or CXCR4, supporting its distinct role in endothelial cells.

**Figure 9.**
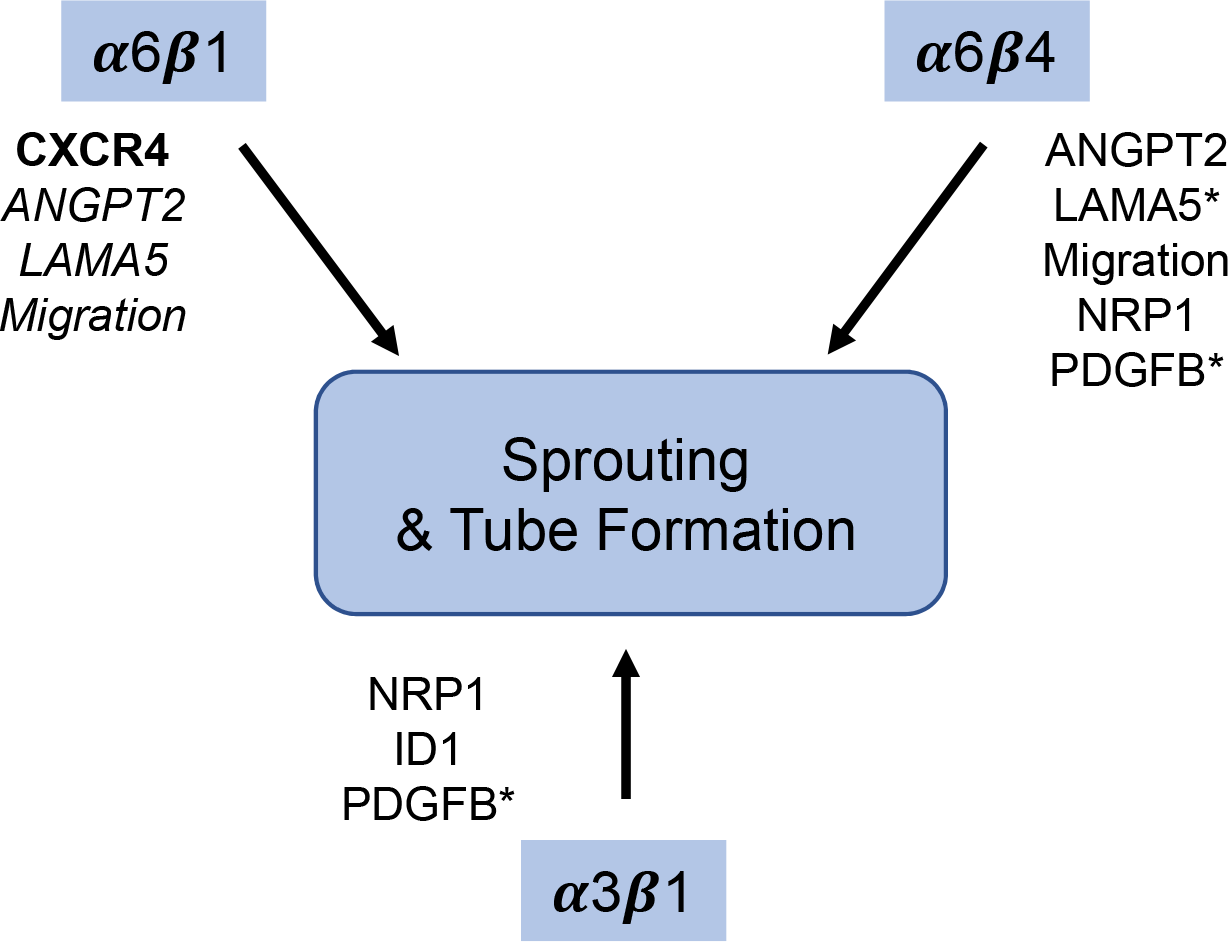
Schematic summary. RNAi-dependent depletion of the α3β1 or the α6 integrins, α6β1 and α6β4 inhibits endothelial sprouting and tube formation in organotypic co-culture models. The expression of CXCR4 shown in bold was inhibited only by the depletion of the integrin α6 subunit and not by depletion of either the β4 or α3 subunit suggesting that α6β1 is the only laminin-binding integrin that impacts CXCR4 expression (Figs 4 & 7 and (Xu et al., 2020)). Depletion of either the α6 or β4 subunit inhibited migration as well as the expression of ANGPT2 and LAMA5 RNA, suggesting that α6β4 regulates the expression of the genes, but does not exclude the possibility of a contribution from α6β1. The depletion of α3β1 or α6β4 inhibited the expression of NRP1 and PDGFB, whereas the expression of ID1 was only inhibited by the depletion of α3β1. *Indicate genes whose expression was significantly inhibited by two of the three siRNA-targeting sequences.

Others have shown that the overexpression of the α6 and β4 subunits in cultured endothelial cells promotes migration on a laminin-332 substrate (Nikolopoulos et al., 2004). We demonstrated that endogenously expressed α6β4 positively regulates migration in cultured endothelial cells without the addition of a laminin substrate or the need for overexpression. Interestingly, our published studies demonstrated that the endothelial expression of laminin-511, a ligand for α6β4, is regulated by α6 integrins (Xu et al., 2020) and Fig. 4C. Thus, α6-dependent regulation of laminin-511 may be involved in promoting migration by α6β4 as observed in our current studies (Fig. 2C) and explain why α3β1 is not required for migration in our assays (Figs. 6D and 7D). It is important to note that others have demonstrated in other cell types that α6β4 can promote migration on non-laminin substrates (O’Connor et al., 1998). Thus, it is possible that α6β4 regulates cell migration by a ligand-independent mechanism.

Consistent with its role in cell migration, we observed the polarized localization of α6β4 during the migratory phase of endothelial tube morphogenesis in planar co-culture. Interestingly, polarized endothelial expression of α6β4 was reported at the tips of endothelial sprouts during neovascularization in human neonatal foreskins (Enenstein and Kramer, 1994). Additionally, previous studies by others have shown that α6β4 promotes epithelial migration and becomes polarized at the leading edge of migrating epithelial cells (Colburn and Jones, 2017; Elaimy et al., 2019; Mercurio et al., 2001). Thus, the α6β4 integrin positively regulates migration in multiple cell types and this regulation seems to involve the specific localization of α6β4.

Others have shown that during the migratory phase of endothelial tube formation in planar co-culture assays, ERK-dependent regulation of cell survival was shown to be critical (Mavria et al., 2006). Multiple different integrin heterodimers regulate ERK activation in many cellular contexts to regulate endothelial cell proliferation and survival (Streuli and Akhtar, 2009). We did not examine ERK activation in our current study; however, our previous studies demonstrated that the loss of α6 integrins did not affect endothelial cell proliferation or survival (Xu et al., 2020). Thus, α6β4 may not significantly contribute to ERK activation, proliferation or survival in our organotypic assays.

Sprouting angiogenesis requires invasion of endothelial cells into the surrounding ECM (Carmeliet and Jain, 2011). The inhibition of sprouting by β4-depleted endothelial cells in our bead-sprout assays suggests that α6β4 may also promote invasion during angiogenesis. Notably, α6β4 has been shown to contribute to cancer cell invasion (Mercurio et al., 2001), indicating that the α6β4 integrin likely contributes to invasion in multiple cellular contexts. Importantly, others have identified a role for endothelial α6β1 integrins both in culture and *in vivo* in the formation of invasive structures known as podosome rosettes, which concentrate the membrane-associated protease, MT1-MMP (Seano et al., 2014). The α6β1, but not the α6β4 integrin was not recruited to rosettes at least when its localization was examined in cell culture (Seano et al., 2014), suggesting that α6β1 and α6β4 likely contribute to invasion by different mechanisms. It will be important to identify the mechanisms by α6β4 contributes to endothelial invasion and how these differ from α6β1.

Our current study identified a role for α6β4 in the regulation of ANGPT2 RNA, although we cannot exclude the possibility that α6β1 also contributes to this regulation. Angiopoietin-2 (product of the ANGPT2 gene) serves as an antagonist to Tie-signaling activated by angiopoietin-1 (Carmeliet and Jain, 2011; del Toro et al., 2010; Huang et al., 2010). Since angiopoietin-1 is secreted by neighboring mural cells *in vivo*, it is unclear how the loss of angiopoietin-2 expression in our organotypic model inhibits morphogenesis. However, some studies have indicated that angiopoietin-2 can activate Tie-2 in an autocrine fashion, suggesting that angiopoietin-2 may have a cell autonomous effect in promoting new vessel formation (Thurston and Daly, 2012). This possibility would be interesting to address in future studies.

Our current results indicate that α6 integrins do not compensate for the loss of α3β1, as depletion of the α3 subunit also inhibited morphogenesis in planar co-culture and bead sprout assays, suggesting that α3β1 regulates these processes by distinct mechanism. Consistent with this notion, the loss of α3β1 expression did not inhibit migration in our assays and affected the expression of a distinct set of angiogenesis-associated genes compared to those regulated by α6 integrins (Supplemental Table 4). *In vivo* studies by da Silva and colleagues (da Silva et al., 2010) reported that endothelial α3β1 acted as a repressor of pathological angiogenesis, as its deletion enhanced tumor associated angiogenesis through the upregulation of KDR (VEGFR2) (da Silva et al., 2010). We did not observe an upregulation of KDR/VEGFR2 at least at the transcript level; however, we did not test whether the post-transcriptional regulation of VEGFR2 was altered in our assay. It is important to note that although da Silva and colleagues demonstrated that the expression of α3β1 was absent from tumor-associated vessels; however, blood vessels in the surrounding normal skin were still positive for α3β1(da Silva et al., 2010). This implies that endothelial α3β1 was present during the initiation of tumor-induced angiogenesis. Thus, this *in vivo* model may not be ideal to examine the role α3β1 during the initial steps of angiogenesis when endothelial cells first respond to angiogenic signals, and may explain the differences with our current findings.

Interestingly, we observed the downregulation of NRP1 and ID1 using all three α3 siRNA targeting sequences. NRP1 is a critical protein enriched in tip cells and functions as a co-receptor for VEGFR2 increasing VEGFR2 signaling to promote angiogenesis (Kofler and Simons, 2015), in part by promoting the formation of filopodia through the activation of CDC42 (Fantin et al., 2015). It will be interesting to determine whether the expression of recombinant NRP1 can rescue the defects in morphogenesis in cells depleted of α3β1. It is important to note that we also observed a downregulation of NRP1 expression using all three siRNAs targeting the β4 subunit, suggesting that also α6β4 contributes to the regulation of NRP1 expression. We did not observe the inhibition of NRP1 expression in α6-depleted cells, presumably because α6β4 is more efficiently depleted when siRNA targeting the β4 subunit employed.

As indicated above, α3β1 promotes the expression of ID1, a transcriptional regulator, which plays a role in angiogenesis during embryogenesis and tumor formation, as well as during endothelial morphogenesis in cell culture models (Benezra, 2001; Benezra et al., 2001; Nishiyama et al., 2005; Tanaka et al., 2010; Volpert et al., 2002). Further studies are needed to determine whether the downregulation of ID1 contributes to the observed phenotype in α3-depleted endothelial cells in our organotypic assays. It is important to note that the α3β1 integrin likely promotes angiogenesis by multiple mechanisms. For example in endothelial cells, α3β1 forms a ternary complex with the tetraspanin CD151 and the matrix metalloprotein and MT1-MMP to promote appropriate proteolysis and the loss of CD151 results in a dramatic loss of α3β1/MT1-MMP association (Yanez-Mo et al., 2008), which has significant implications as CD151 has been shown to promote angiogenesis (Sadej et al., 2014). Interestingly, α3β1 has been associated with tumor cell invasion (Mitchell et al., 2010) and has been shown to regulate MMP-9 RNA stability in keratinocytes (DiPersio et al., 2000; Iyer et al., 2005). Given the matrix-dense environment of the two organotypic assays employed in our studies, the positive regulation of proteases by endothelial α3β1 during morphogenesis could possibly explain the lack of tube formation and sprouting by α3-depleted endothelial cells in planar co-cultures and bead-sprout assays, respectively.

In summary, our current studies demonstrate that the expression of α6β4 and α3β1 is required for endothelial sprouting and tube formation in our two organotypic angiogenesis assays. Our previous studies showed that the expression of α6 integrins is required to maintain the morphological integrity of endothelial tubes, as well as the expression/association of laminin 511 with these tubes (Xu et al., 2020). We have yet to determine whether this phenotype requires the expression of α6β1 and/or α6β4. However, it is important to note that the endothelial specific knockout of the β4 subunit (ITGB4) or the α5 chain of laminin-511 resulted in defective barrier function in the neural microvasculature (Song et al., 2017; Welser et al., 2017), indicating that α6β4 can play a role in maintaining endothelial tube integrity. Our current study suggests this may be specific for α6 integrins, as depletion of α3β1 from established tubes did not alter tube morphology or the expression laminin-511. Finally, the organotypic assays employed in our studies will provide useful reductionist systems to dissect the molecular mechanisms by which individual laminin-binding integrin contribute to endothelial morphogenesis.

## Materials and Methods

### Cell Culture

Human umbilical vein endothelial cells (HUVECs) were from Lonza (Allendale, NJ) and were cultured in in EGM-2 (Lonza, CC-3162) and used between passages 2 - 6. Adult human dermal fibroblasts (HDFs) were isolated and characterized as previously described (Varney et al., 2016; Zheng et al., 2019) and generously provided by the Van De Water laboratory (Albany Medical College) and used between passages 8 - 14. Human embryonic kidney epithelial 293FT cells (HEK293FT) were a kind gift from Dr. Alejandro Pablo Adam lab (AMC). Cell lines were authenticated and routinely tested for contamination. HDFs and HEK293FT cells were cultured in DMEM (Sigma D6429) containing 10% FBS (Atlanta Biologicals), 100 units/ml penicillin (Life Technologies), 100 µg/ml streptomycin (Life Technologies), and 2.92 µg/ml L-glutamine (GE LifeSciences). All cells were cultured at 37°C in 5% CO2.

### siRNA

HUVECs were plated in 6-well tissue culture plates and transfected with siRNA at a 50 nM concentration with RNAiMAX (ThermoFisher) using the protocol provider by the manufacturer. HUVECs transfected with siRNA were assayed for knockdown and used in planar co-cultures (described below) 48 h after transfection. HUVECs transfected with siRNA and used in bead sprout assays (described below) were transfected during bead coating and assayed for knockdown at the end of experiment. Sources and nucleotide sequences of siRNAs used in this study are provide in Table S2.

### Inducible shRNA

Doxycycline-inducible lentiviral (SMART) vectors harboring shRNAs targeting the α3 integrin subunit or a non-targeting (NT) shRNA were purchased from Dharmacon (Lafayette, CO). Lentiviruses were produced by co-transfection of HEK293FT cells with the shRNA expression vector together with the packaging plasmid, psPAX2, coding for Gag, Pol, Rev, Tat (#12260, Addgene), and the envelope plasmid, pMD2.G, coding for VSV-G (#12259, Addgene). HUVECs were transduced with filtered viral supernatant plus 8 µg/ml polybrene. Cells were induced with doxycycline (100 ng/ml) for 48 h prior to adding HUVECs to co-cultures or in some experiments after 10 days in co-culture as described below. Nucleotide sequences of shRNAs of shRNAs used in this study are provided in Table S2.

### Quantitative PCR (qPCR)

TRIzol (ThermoFisher) was used to isolate RNA from siRNA transfected HUVECs, as well as, HUVECs expressing shRNAs. Extraction of RNA from bead sprout assays (described below) using TRIzol was performed after the removal of HDFs with trypsin-EDTA Solution 10X (59418C, Sigma). cDNA was synthesized with iScript Reverse Transcription Supermix (BioRad) using 1 µg of RNA. Equal amounts of cDNA were used in qPCR reactions performed with iQ SYBR Green Supermix (BioRad). The nucleotide sequences of the qPCR primers used in this study are listed in Table S3.

### Migration

SMART Vector shRNA expressing cells were induced with doxycycline (100 ng/ml) for 48 h prior to assays. Cells were cultured in serum-free EGM-2 medium overnight prior to assay. Similarly, siRNA transfected cells were serum starved 48 h post transfection. Fifty thousand cells were seeded in triplicates into transwells and a separate 24-well plate. Serum-containing EGM-2 was then added to the lower chamber of transwells followed by incubation for 4 hours at 37°C in 5% CO2. Transwells were fixed with 4% PFA (Electron Microscopy Sciences) and stained with DAPI. The lower membrane was imaged with a 4X objective and density quantified using ImageJ (NIH). Cell seeding efficiency was determined by performing toluidine blue assays in the 24-well plates. These assays were performed by fixing cells with 70% ethanol at room temperature for 1 hour, followed by wash with dH2O and staining with 0.05% toluidine blue at room temperature for an additional 2 hours. After wash with dH2O, toluidine blue was extracted with 10% acetic acid at 0.3 ml / well and absorbance measured at 650 nm, using 405 nm as reference on a Synergy2 microplate reader (BioTek Instruments). An empty well was processed the same way and used for baseline. Migration efficiency was determined by dividing DAPI density by absorbance.

### Western blotting

Western blotting was used to confirm RNAi induced knockdown. Cells were lysed in mRIPA buffer (50 mM Tris pH 7.4, 1% NP-40, 0.25% Na Deoxycholate, 150 mM NaCl, 1 mM EDTA) containing both phosphatase (Sigma, #4906837001) and protease inhibitor cocktails (ThermoFisher, 78440). Equal amounts of protein (20 to 40 µg) were separated by SDS-PAGE and transferred to nitrocellulose for antibody probing. Imaging was performed with a ChemiDoc XRS+ (BioRad) and quantitation with Image Lab (BioRad).

### Organotypic culture assays

#### Planar co-culture

As one organotypic culture, we utilized the planar co-culture model developed by Bishop and colleagues (Bishop et al., 1999) and modified by the Pumiglia lab (Bajaj et al., 2012). This model reconstitutes some of the complex interactions that occur during angiogenesis among endothelial cells, the ECM and supporting cells. To set up the co-culture, HDFs were seeded in tissue cultures dishes ± glass coverslips and cultured to confluence. The medium was changed to EGM-2. HUVECs, expressing targeting or non-targeting RNAi, were then seeded 16 h later at a density of 20,000 cells per 9 cm^2^ and cultured up to 10 days. ShRNA-mediated knockdown in pre-formed tubes was accomplished by culturing HUVECs expressing doxycycline-inducible lentiviral (SMART) vectors on HDFs for 10 days, in the absence of doxycycline with medium changed every 48 h. Doxycycline was then introduced on day 11, at a concentration of 100 ng/ml and refreshed every 48 h for up to 16 days. Endothelial morphogenesis and changes in tube structure were analyzed by immunofluorescence microscopy.

#### Bead-Sprout assay

To study endothelial sprouting, we employed the bead sprout assay as described by Nakatsu and Hughes (Nakatsu and Hughes, 2008). Cytodex 3 beads (GE) were coated at ∼1000 HUVECs per bead inside of a 2 ml microcentrifuge tube for 4 at 37°C, mixing gently by inverting the tubes every 20 min and transferred to a T25 flask and incubated at 37°C, overnight. Beads were then washed 3X with EGM-2 medium and re-suspended in PBS containing 3 mg/ml of fibrinogen (Sigma, #F8630) and 0.15 U/ml of aprotinin (Sigma, #A6279). Thrombin (Sigma, #T4648) was added at a final concentration of 0.125 U/ml and the mixture was plated in wells of and 8-well slide (Corning, #3-35411). The mixture was allowed to clot for 30 min at 37°C. HDFs were then added to the top surface of the fibrin gel in EGM-2 medium at a concentration of 30,000 cells per well. The formation of sprouts and sprout lengths were assayed by either immunofluorescence of phase contrast microscopy.

### Immunofluorescence Microscopy

Planar co-cultures were fixed with 4% PFA (Electron Microscopy Sciences) for 15 min, permeabilized with 0.5% Triton X-100 in PBS for 15 min, and then blocked with 2% BSA in PBST for 1 h at RT. Antibodies were diluted in 2% BSA in PBST and incubated with cells overnight at 4°C. Samples were then washed 4X with PBST at RT over the course of 4 h, and then incubated for 1 h with the appropriate secondary antibodies (1:1000 dilution). Following secondary antibody staining, samples were washed 3X with PBST at RT for 1 h and mounted with SlowFade Gold antifade reagent (ThermoFisher). Samples were analyzed using a Nikon inverted TE2000-E microscope equipped with phase contrast and epifluorescence, a digital CoolSNAP HQ camera, a Prior ProScanII motorized stage and a Nikon C1 confocal system and EZC1 and NIS-Elements acquisition software. Images were acquired with Plan Fluor 4X/0.13, Plan Fluor 10X/0.30, Plan Fluor ELWD 20X/0.45, Plan Apo 40X/1.0 oil, and Plan Apo 100X/1.4 oil objectives and analyzed with either NIS elements (Nikon). Contrast and/or brightness were adjusted for some images to assist in visualization. Fibroblasts were removed from bead sprout assays using trypsin-EDTA Solution 10X (59418C, Sigma) prior to fixation and staining. See Table S1 for antibody information.

### Image analysis

For planar co-cultures, where endothelial cells initially form cell trains/cord, which mature into endothelial tubes, cord/tube lengths in planar co-cultures were measured by tracing tubes within each field using NIS Elements (Nikon). Any tubes that extend beyond the field were excluded from analysis. Tube widths in planar co-cultures were calculated with AngioTool (NIH) by dividing total tube area by total tube lengths per field. Average tube length per field was calculated with AngioTool (NIH). Sprout lengths in bead sprout assays were measured by tracing each sprout using NIS elements (Nikon) and sprouts per bead were counted manually.

### Western blotting

Western blot analysis was used to assay the expression of the integrin α3 subunit and integrin β4 subunit. Cells were lysed in mRIPA buffer containing a protease/phosphatase inhibitor cocktail used at a 1:100 dilution. Equal volumes of protein were separated by 10% SDS-PAGE and transferred onto either 0.4 mm nitrocellulose membranes for analysis. See Table S1 for antibody information.

### Animal experiments

All animal experiments and procedures were performed in accordance with the Albany Medical College Institutional Animal Care and Use Committee (IACUC) regulations. In accordance with protocols approved by the Albany Medical College IACUC. Adult murine retinas and ears were harvested from both male and female C57BL/6 mice between the ages of 8 – 12 weeks. Adult murine skins were harvested from male and female C57BL/6 mice expressing the transgenes: LSL-tdTomato and VE-Cadherin driven CreERT2. Cre recombinase activity was induced with via oral gavage of 10 µg/g of tamoxifen per day for 5 days.

### Statistical analysis

Statistical analysis was performed with GraphPad Prism software using Student’s t-test or one-way ANOVA with post-hoc analysis using Dunnett’s or Tukey’s multiple comparisons tests as indicated in the Figure Legends. P value of p<0.05 was considered to be statistically significant.

## Acknowledgements

The authors thank Drs. C. Michael DiPersio and Livingston Van De Water for critically reading our manuscript and for their helpful comments, Dr. Livingston Van De Water for providing human adult dermal fibroblasts, Debbie Moran for assistance in the preparation of the figures.

---

**Table S1:**
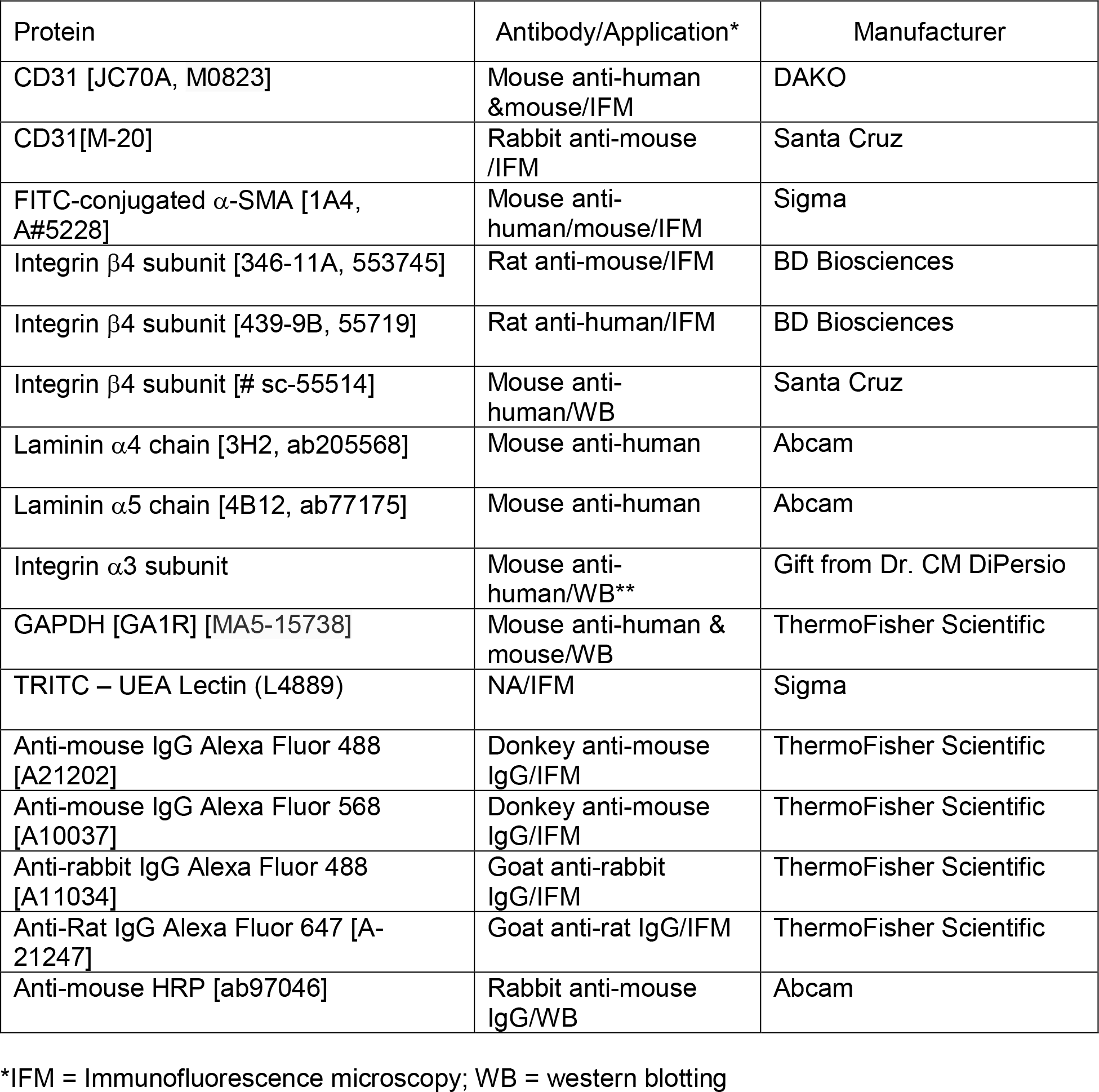
Antibodies

**Table S2:**
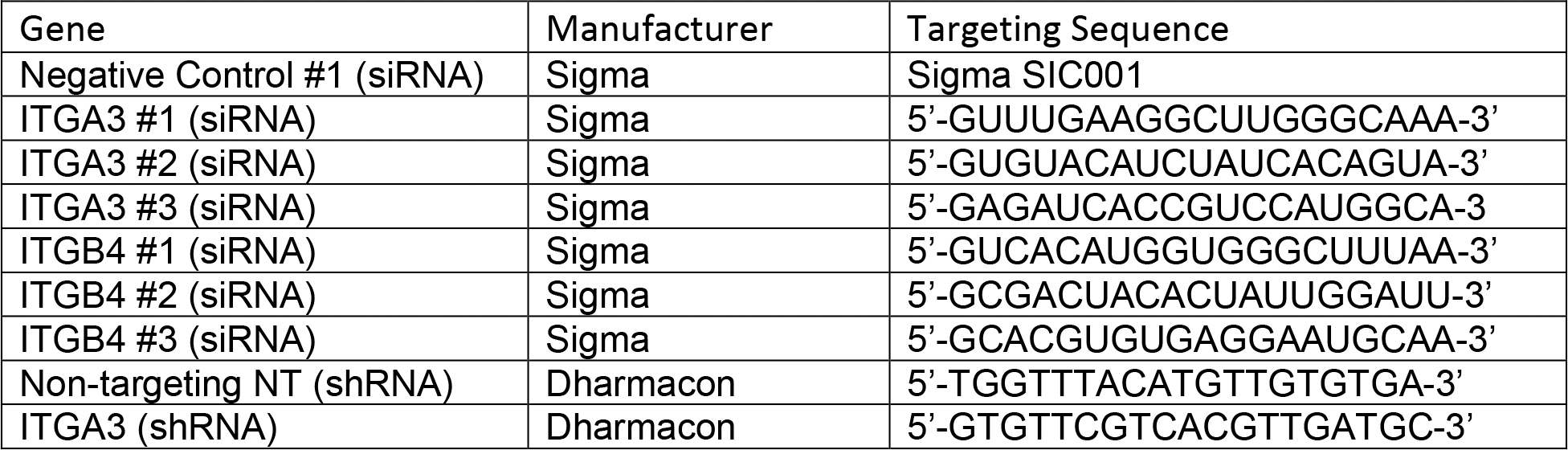
RNAi

**Table S3:**
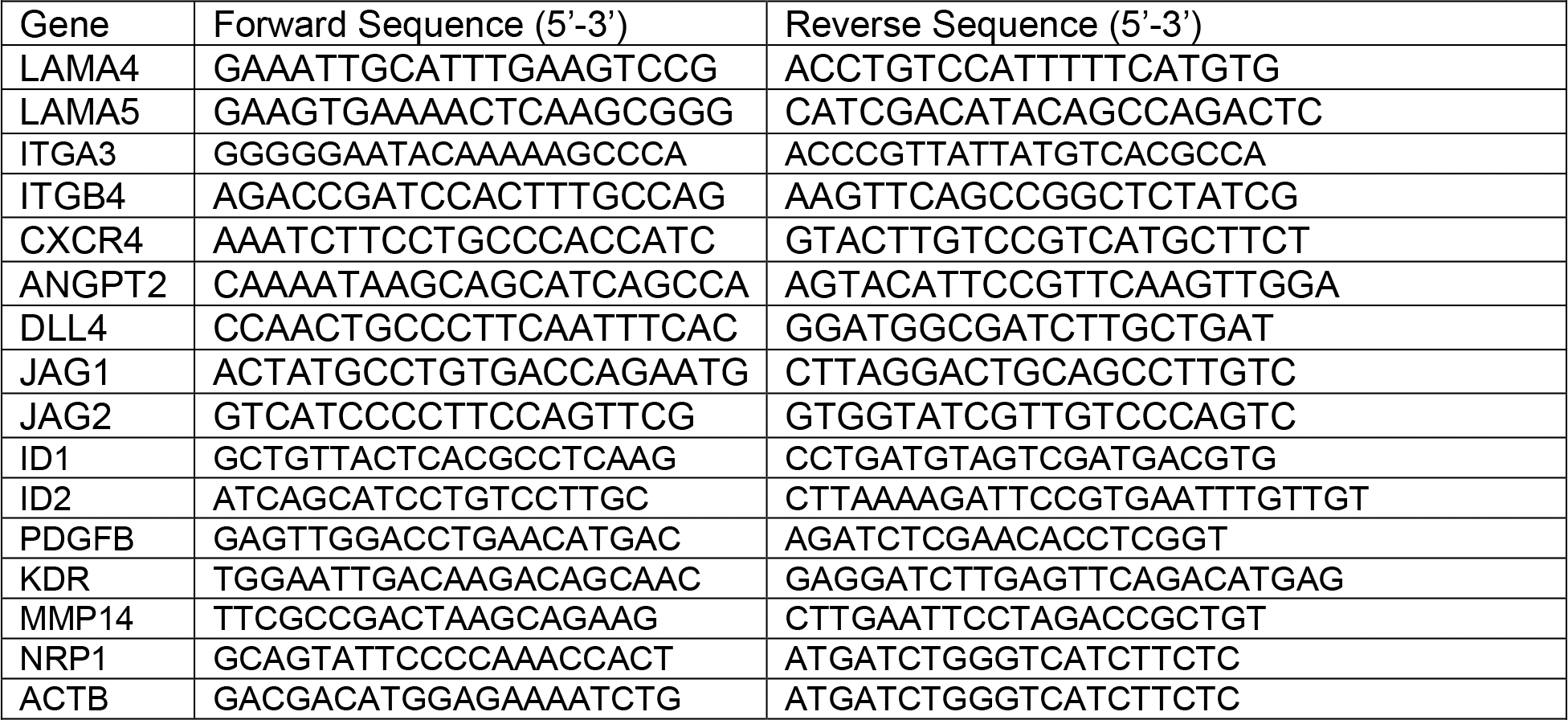
QPCR Primers

**Table S4:**
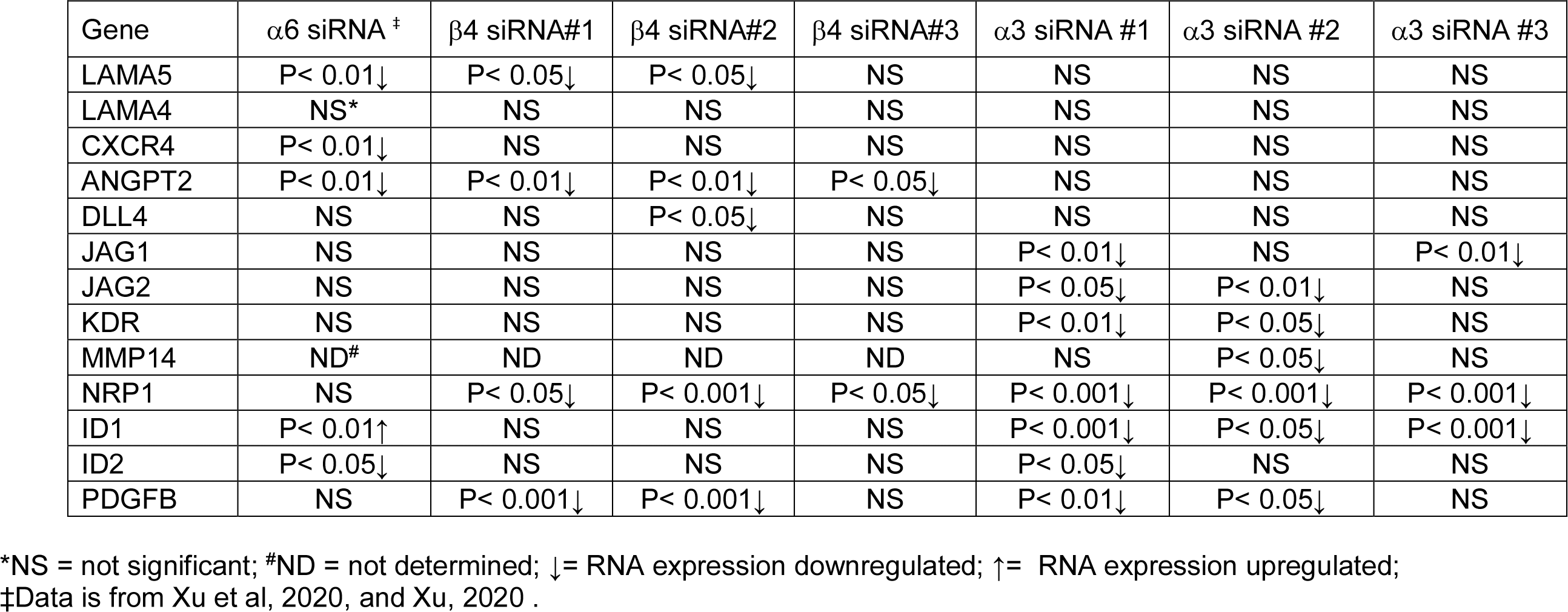
Summary of Gene Expression Analysis

| Gene | $\alpha 6$ siRNA <sup>†</sup> | $\beta 4$ siRNA#1 | $\beta 4$ siRNA#2 | $\beta 4$ siRNA#3 | $\alpha 3$ siRNA #1 | $\alpha 3$ siRNA #2 | $\alpha 3$ siRNA #3 |
| --- | --- | --- | --- | --- | --- | --- | --- |
| LAMA5 | P < 0.01↓ | P < 0.05↓ | P < 0.05↓ | NS | NS | NS | NS |
| LAMA4 | NS* | NS | NS | NS | NS | NS | NS |
| CXCR4 | P < 0.01↓ | NS | NS | NS | NS | NS | NS |
| ANGPT2 | P < 0.01↓ | P < 0.01↓ | P < 0.01↓ | P < 0.05↓ | NS | NS | NS |
| DLL4 | NS | NS | P < 0.05↓ | NS | NS | NS | NS |
| JAG1 | NS | NS | NS | NS | P < 0.01↓ | NS | P < 0.01↓ |
| JAG2 | NS | NS | NS | NS | P < 0.05↓ | P < 0.01↓ | NS |
| KDR | NS | NS | NS | NS | P < 0.01↓ | P < 0.05↓ | NS |
| MMP14 | ND<sup>#</sup> | ND | ND | ND | NS | P < 0.05↓ | NS |
| NRP1 | NS | P < 0.05↓ | P < 0.001↓ | P < 0.05↓ | P < 0.001↓ | P < 0.001↓ | P < 0.001↓ |
| ID1 | P < 0.01↑ | NS | NS | NS | P < 0.001↓ | P < 0.05↓ | P < 0.001↓ |
| ID2 | P < 0.05↓ | NS | NS | NS | P < 0.05↓ | NS | NS |
| PDGFB | NS | P < 0.001↓ | P < 0.001↓ | NS | P < 0.01↓ | P < 0.05↓ | NS |

\*NS = not significant; <sup>#</sup>ND = not determined; ↓ = RNA expression downregulated; ↑ = RNA expression upregulated;
†Data is from Xu et al, 2020, and Xu, 2020 .

**Supplementary Figure 1.**
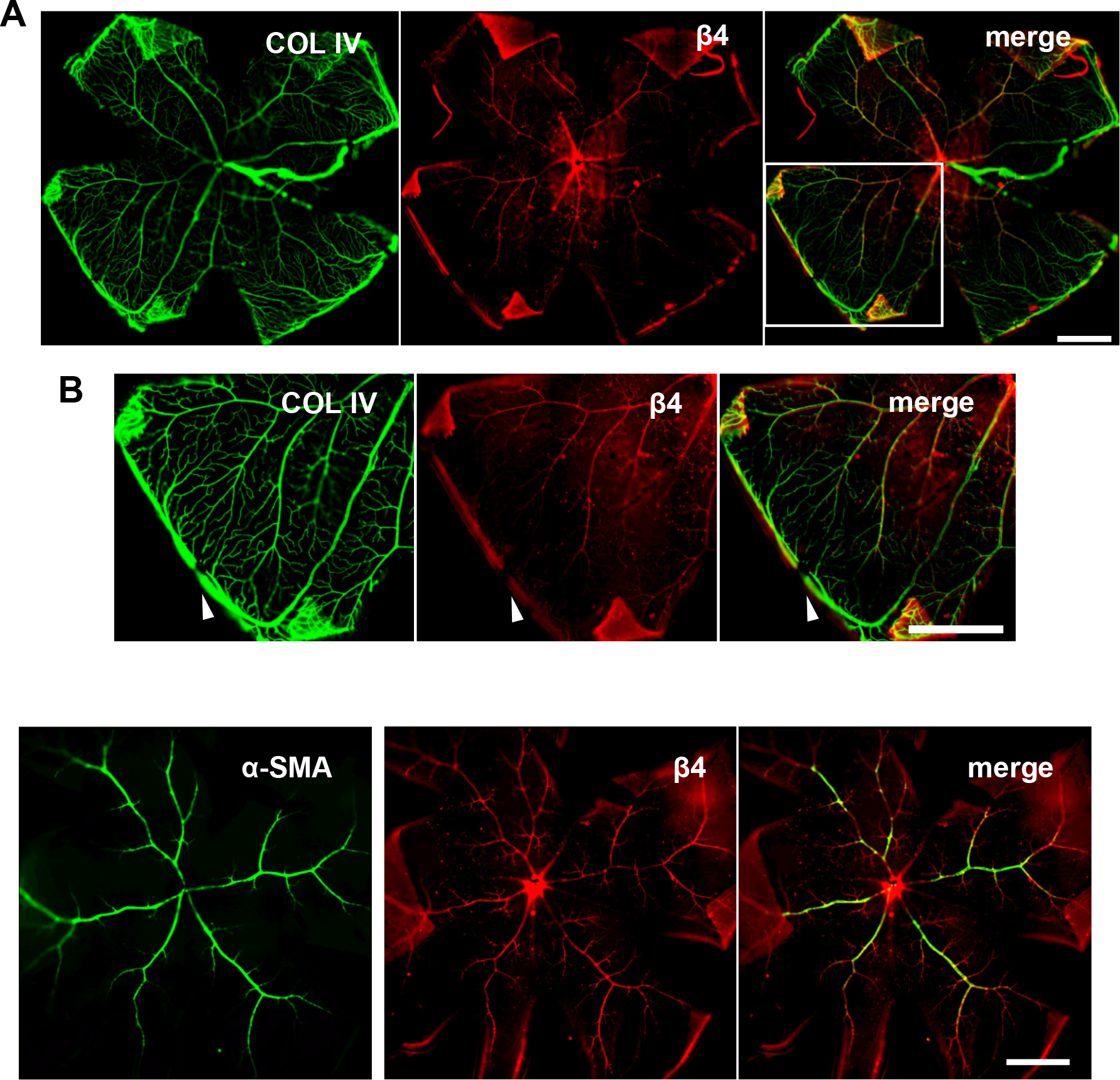
Integrin α6β4 is primarily expressed in the arterioles of adult mouse retina. **(A-B)** Integrin α6β4 is expressed in a subset of vessels of adult mouse retina. **(A)** Immunofluorescence staining of whole mount mouse retina for Collagen IV and β4-integrin subunit. **(B)** Enlarged image of one petal of retina showing restricted β4 expression. Arrowhead and arrows showing β4-positive and negative vessels, respectively. Scale = 1 mm. **(C)** Immunofluorescence staining of whole mount mouse retina for α-smooth muscle actin (α-SMA) and β4-integrin subunit. Arrow heads show examples α-SMA / β4 co-localization in the arterial branch. of Scale = 1 mm.

**Supplementary Figure 2.**
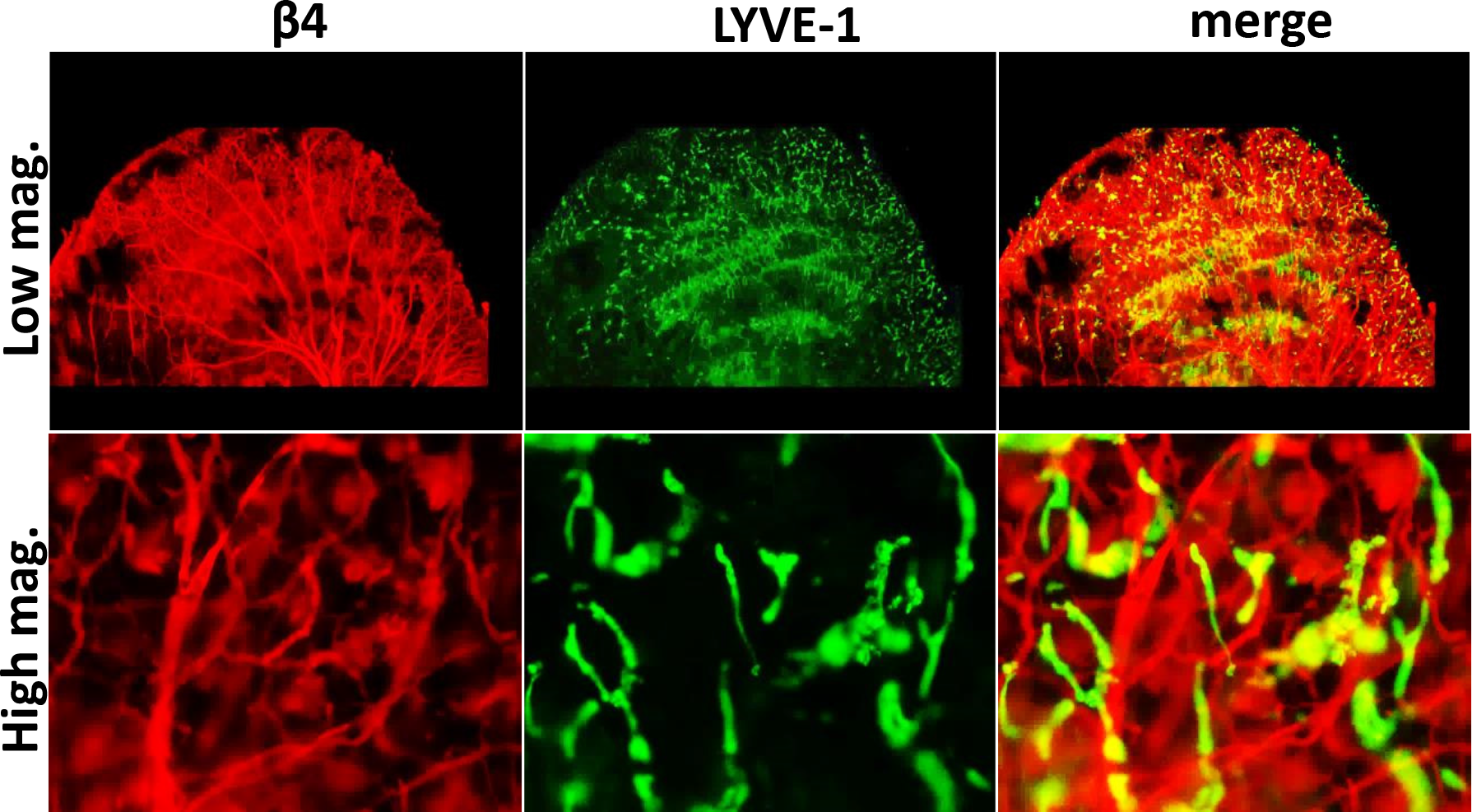
Integrin α6β4 is not expressed by the lymphatic system of the adult mouse dermis. Immunofluorescence staining of whole mount mouse ear for lymphatic <u>v</u>essel <u>e</u>ndothelial <u>r</u>eceptor 1 (LYVE-1) and β4-integrin subunit.

**Supplemental Figure 3.**
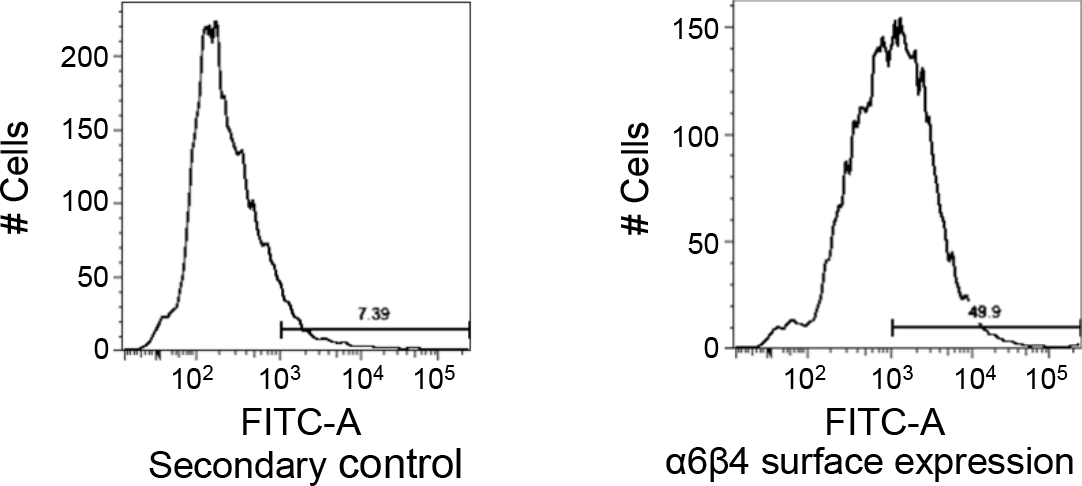
Flow cytometry was used to analyze surface expression of integrin α6β4 on human umbilical endothelial cells (HUVECs).

